# COULD PLANTS BE SENTIENT?

**DOI:** 10.1101/121731

**Authors:** Paco Calvo, Vaidurya Sahi, Anthony Trewavas

## Abstract

Feelings in humans are mental states representing groups of physiological functions that usually have defined behavioural objectives or purpose. Feelings are thought to be coordinated in the brain stem of animals and are evolutionarily ancient. One function of the brain is to prioritise between competing mental states, and thus groups of physiological functions and in turn behaviour. Anger, fear or pain call for immediate action whereas hunger, or thirst, signify longer term needs and a requirement for search. Plants use groups of coordinated physiological activities to deal with defined environmental situations but currently have no known mental state to prioritise any order of response. Plants do have a nervous system based on phloem which is highly cross linked. Its potential for forming a mental state is unknown but it could be used to distinguish between different and even contradictory signals and thus determine a priority of response. The vascular nervous system stretches throughout the whole plant providing the potential for assessment in all parts and commensurate with its self-organising, phenotypically plastic behaviour.

## INTRODUCTION

Sentience is commonly regarded as the capacity to feel subjectively and is used to distinguish feelings from reason or logic. Feelings are mental experiences of body states. But they are subjective making it difficult to know if the internal experience is even identical between different human individuals. Whether animals are sentient is a question that gives rise to huge controversy (Boyle, 2009). Could plants be sentient? Even if the evidence was encouraging, it would always remain unanswerable.

Sentience is generally considered limited to organisms that have a nervous system and a centralised brain. For organisms without these supposed requirements, the notion of sentience has been rejected out of hand. Plants are placed in this category (Grinde, 2013. Animal Ethics Inc. www.animal-ethics.org/beings-conscious). The reasons for rejection are four fold

The supposed absence of a mechanism for transmission of information similar to the animal nervous system. This article shows that plants do have such a mechanism, something known a century ago.
Plants don’t have brains the supposed seat of feelings. Again there is reason to doubt this claim as will be described later.
Plants are simple. They don’t move and thus don’t need a nervous system. A recent attempt to compare the complexity of large angiosperms with large animals using discrete complexity criteria failed to distinguish the two (Trewavas, 2014). Long range communication within any plant is essential to balanced development, growth and survival.
The capacity to feel arose in evolutionary terms solely from its usefulness in motivating animals; it doesn’t make sense for plants that can’t run away from a threat or forage for a food they enjoy.

On the contrary threats to plant life are very common and are counteracted. Aspects of the behaviour of some like parasitic plants are clearly motivated and they forage sensibly (Trewavas, 2017). This article deals with these reasons for rejection and points to errors in knowledge which are not uncommon amongst many animal scientists and even some who research plants (Chamovitz, 2012).

### Perceptual bias generates erroneous understanding of plant behaviour

Subjective, anthropomorphic attitudes are commonly used to judge plant behaviour but the bias often goes unrecognised. Behaviour is classed as the specific response to one or a group of stimuli. In animals it is easy to characterise because it usually results in visible movement. Absence of visible movement leads to common assessments that plants lack behaviour altogether with, of course, a few exceptions like Dionaea, (Venus fly trap) or the sensitive plant (*Mimosa pudica*). However our ability to see movement is constrained within discrete limits (Trewavas, 2014). The perceptual bias involved can be illustrated by asking instead what is the commonest form of biological behaviour.

Amongst virtually all angiosperm plants, behaviour can be characterised as either molecular changes in composition, or reversible movements of leaves, tendrils, stomatal cells for example, or the more easily seen but slower phenotypic plasticity. It is not difficult to see the plasticity of shoots when looking at the branching patterns of two deciduous trees of the same species and age and noting, when present, how the positions and characteristics of branching reflects close-by neighbours. Light is the source of energy for virtually all plants (some 5000 species of parasitic plants being exceptions) and is fought for when neighbours are sufficiently close to compete. Equally complex competitive behaviour is exhibited by the root system something rarely seen except under exceptional experimental conditions. Long range communication is essential to ensure reasonable balance between shoot (trunk) and root. Time lapse photography of the shoot has fortunately started to change perspectives on plant behaviour and there are many examples on youtube. The phenotype is a history of the environmental conditions it has experienced.

99% or more of eucaryotic life on this planet is however plant, not animal as indicated by the ratio of oxygen to carbon dioxide. Thus the commonest form of behaviour is not movement we see but that expressed by plants. Phenotypic plasticity certainly occurs in animals and humans too. Weight lifters and all those obsessed with the manipulation of the body beautiful are cases in point. Again note that these changes which, in wild circumstances, would reflect on fitness, are only detectable after weeks to months; similar to plant plasticity.

### These differences in behaviour probably started with the first eukaryotic cells

Animals and plants separated in evolutionary terms at probably the single cell stage. The fundamental difference lies in the means to acquire necessary energy for survival. Movement was nearly always essential for primordial animal cells to find food. Predator-prey relationships amongst animals, almost certainly accelerated the familiar characteristics of elaborate sensory systems coupled with a highly evolved musculature. Finally these were coupled together by a fast communication nervous channel, that increased the probability of the predator to capture prey or for the prey to escape being eaten.

The primordial plant cell on the other hand, acquired a blue green algal symbiont and became photosynthetic. Photosynthetic products are often osmotically active. To prevent these photosynthetic cells from exploding, it was necessary to concomitantly evolve a relatively rigid wall, a severe impediment to movement. Coupled with the ubiquity of light around the earth, evolutionary pressure to move quickly and easily was never an evolutionary imperative.

### Plant behaviour is complex not simple

Plants and animals are entirely different eukaryotic solutions to the problems of survival. But there are marked differences in perception. Roving animals in search of prey need only make a cursory search of any environment. Even insect herbivores need only detect green vegetation for feeding.

Being largely confined to the place in which they germinate and grow, higher plants distinguish the features of their environment with much greater discrimination. The resources necessary for life, minerals, water, light, are effective plant food. But they are patchily distributed around the plant body and are fought over competitively not only to acquire them but to acquire them first and deny them to others. Fine discrimination leads to better exploitation but an exploitation that requires growth and the internal resources from either root or shoot to enable plasticity in growth. In those plants that have been investigated there is competition for these internal resources too. Decisions have therefore to be made out of a range of possibilities; those plants that can make them more quickly, with lower cost are more likely to accumulate more food, produce more seeds and siblings and thus are fitter (Trewavas, 2014; 2016). Nature red in tooth and claw might describe the animal world but in the plant world it is green, overgrowth and phenotypic plasticity.

### Threats to survival of the individual plant

Herbivory, other physical damage by for example wind or trampling, disease, light or shade, drought, flooding, cold and hot temperature extremes all initiate dramatic phenotypic responses including resistance unless overwhelmed. Less stressful episodes of cold, heat, water loss or mechanical stress by wind, are learnt so that further episodes are dealt with more robustly, more quickly and with greatly improved resistance to the threat. Herbivory and disease prime the plant. After the first episode of insect damage, any subsequent attacks again are responded to more rapidly and greatly enhanced in size. Priming is clearly a learning mechanism and the induced memory lasts for years and in some plants survives reproduction (Frost et al., 2008; Zimmermann et al., 2016). Basically it is behaviour profiting from experience, a common definition of intelligence (Trewavas, 2017). Changes in chromatin structure are its likely basis (Ali et al., 2013). Recognition of the species of attacking caterpillar is gained through recognition of the precise chemical composition of salivary juice. In turn the attacked plant emits complex signatures, mixtures of volatile chemicals that are recognised by parasitoids of the specific caterpillar, a kind of burglar alarm with associated police response (Diezel et al., 2009). Further attacks lead to a much quicker and greatly elevated resistance response. Priming can last years and even in some cases survives reproduction.

Similar problems beset the root. Soil strength, stones, compaction, all require morphological change and phenotypic adaptation; disease and nematode damage require a plethora of responses. Interactions with various fungal species can lead to beneficial symbiosis ranging all the way down to parasiticism or disease. Roots from adjacent competitive neighbours are sensed and lead to proliferation to deny resources to the neighbour as well as the benefits of acquiring the soil resources first. A variety of soil chemicals are sensed and acted upon. Each signal constructs a different phenotype with a unique underlying molecular base. Mechanisms that transduce many of these environmental signals are known (Trewavas, 2014). Errors in behavioural responses are recognised and corrected (Trewavas, 2017). There is nothing simple here.

Differing groups of signals appear at different times and can be contradictory. A key question then is how plants place weight upon any of the signals perceived and how they prioritise them in terms of response. That is one of the questions this article attempts to raise.

Growing wild plants self-organise; each stage of development acts as a platform for the next. Development itself is thus a learning process and the unpredictable environment acts as the context in which learning takes place. Higher plant organisation can be described as analogous to a republic in contrast to the monarchical animal. However the organisation of a republic is still coherent but permits a much greater level of distributed and local control than is present in a monarchical or dictatorial system. These two great kingdoms reflect different facets of organisation that in turn reflect the different evolutionary decisions made when both were single cells as to the means for acquiring energy. Animal development is constrained within relatively rigid boundaries; that is the price paid for the lifestyle that necessitates movement.

## THE NATURE OF FEELINGS IN ANIMALS

Sentience is generally held to describe the presence of emotions and feelings in human beings as contrasted with logic and reason. Whilst humans can put words to internal experiences such as hunger, pain, fear, anger, well-being or the more nuanced feelings of compassion, gratitude, even sexual love these are, and recognisably, entirely subjective even to the human individual (Damasio and Carvalho, 2013). Their presence in ourselves however has given rise to an enormous discussion and literature arguing for equivalent feelings in other animals (Boyle, 2003). The best that can be suggested, usually in discussion of pain, is a kind of insurance; we should avoid certain actions with animals, in case it is indeed equivalent in intensity and damage to that in human beings.

Charles Darwin (1872) in his book on animal emotions extensively summarised previous studies and combined them with his own considerable observations most notably on zoo primates. He also included cats, dogs, horses etc., in his discussion. By observing behaviours which he could interpret as analogous to those in humans, he considered that emotions were present. Anthropocentric attitudes again complicate the issue of assessing primate behaviour, experience and emotion. Two well-known primates, Washoe and Koko, chimpanzee and gorilla respectively and others, were taught sign language and acquired some 300-1000 words after several years of training (Gardener et al., 1989; Patterson and Linden 1981). Intellectually these primates were rated as equivalent to a 2-3 year old human child. But how well would a 2-3 year old human child survive in the wild circumstances familiar to chimps and gorillas which they manage with alacrity? Chimps recognise in a few seconds their social position when placed in a new tribe, a behaviour crucial to their survival. What emotion is involved there? Is it well-being, fear, or a simple survival mechanism performed without any emotional content?

There is more to intellect than learning signs for language and trying to identify apparent emotions. Human language is complex and in its complexity maybe unique, but judgements made on language alone are entirely flawed. In their true wild context these animals are surely highly intelligent and their survival depends upon it.

Strong drivers of feelings are the nociceptors. These transmit information to the brain on tissue damage and the detection of noxious or potentially noxious circumstances eliciting the sensation of pain. Whether a nervous system is essential for sensing and response to damage can be queried. Jennings (1926) discusses sentience, experience and cognition in the context of single-celled animals. When *Amoeba, Stentor* or *Paramecium* perceive localised, damaging circumstances they move in the opposite direction and into what, we judge to be, a more equitable environment or one in which threat is diminished. Is this behaviour he asked, consistent with what would be expected if the single cell experienced the equivalent of pain in noxious circumstances? He concluded that it would, but added the obvious proviso that it could never be known if pain was involved. Are then nervous systems really a prerequisite for feelings? Or are they merely an elaboration of sensory experience, feelings experienced even by the simplest of organisms?

As to the actual human experience of pain, it is difficult to assess how well we distinguish actual pain from our perception that it must be painful. There are certainly a range of sensitivities between different humans from highly sensitive to effectively null experience of pain itself. There are, as well, good anecdotal examples of men who lost limbs in battle with seemingly little painful effect because they continued to fight. Animals do react when injured in a way that we would interpret as painful to us. But can we ever know their actual experience?

### Human feelings have an evolutionary benefit

Feelings in humans like all other human characteristics are present because they served a role in natural selection and subsequent evolution. They represent a mental state which is connected to groups of physiological and metabolic activities focussed on required individual behaviours (Damasio and Carvalho 2013). Perhaps the most familiar to the reader is that of flight or fight, which can vary enormously in intensity between human individuals. The threat signal generates a mental state involved in energising the familiar group of physiological responses; increased cardiac and respiratory activity, elevated blood flow rates, blood sugar, dilated pupil and increased secretions of adrenalin and cortisol amongst others. By providing the necessary assessment of a potential or potentially threatening future, the brain constructs priorities between competing problems. Amongst a plethora of potential behaviours, which one needs attention first.

Behaviours in animals as in plants are a way of maintaining as far as possible what might be termed a state of life cycle well-being. It enables the individual to continue the path to reproduction with least interruption and culminating in satisfactory reproduction. It is not clear that well-being is a definable mental state in humans. But well-being circumstances may be easier to characterise for many crop plants. Good, well-drained soil with nutriment and water readily available, no wind, an optimal temperature, freedom from disease and herbivory and an abundance of pollinators. Reproductive abundance is the consequence, a commonly-used proxy in the wild for fitness. But these crop plants have been selected from the enormous range of genotypes in the wild and bred to respond well to these circumstances.

Feelings are thought to originate in the brain stem first and thus their evolution is probably ancient (Damasio and Carvalho, 2013). What came first was the grouping of physiological responses together in response to defined environmental perturbations; only later it is thought were these coordinated by nervous activity. The incorporation of mental states helped provide the organism with a potential guide to adaptive behaviours including intelligent learning and memory. Gardner (1983) in his creative approach to intelligence, incorporates both emotional and social intelligence as critical aspects of this behaviour. An arguable case can be made for single cells expressing many of the fight or flight biochemical responses in response to a damaging environment.

### Groups of plant physiological and phenotypic changes are induced by specific signals

There is organisational similarity in the way that both animals and plants respond to external signals. Perception leads to assessment and in turn a sensible response. In higher animals, perception involves detection in one or more sensory systems and is communicated to assessment areas in the brain via nervous connections or hormones. Responses are initiated through activating specific muscles and coincident usually with new hormone release. Furthermore the organism monitors whether the response is sufficient leading to further action if it isn’t. The immune system, which both learns and remembers, may be the exception in not directly using nervous communication but its behaviour is generally integrated with the whole organism.

Perception in plants is located in groups of cells, whole tissue or tissues. There are subsequent downstream effects that often involve long distance communication to other tissues or organs. Chemical communication is common and complex, involving small and large RNAs, proteins, peptides, oligosaccharides and a large range of smaller chemicals some of which are designated as hormones although these usually lack the discrete site of synthesis in a gland or tissue familiar with mammalian endocrinology (Trewavas, 2014). Even minerals and water can be employed to communicate internal information. Finally there is electrical communication which follows on this section of the article.

At some stage the nature of the perceived signals are assessed and subsequent downstream changes initiated leading for example redirection of leaf position, closure of leaf stomata, or tendril curling taking some 10-15 minutes. Phenotypic changes obviously take much longer and probably use a variety of feedback mechanisms to optimise the appearance and growth of the new structure. There is evidence that predictions are commonly made of potential futures and included in the eventual responses. One feature of plant response currently lacking is the identity of signals that indicates to remote tissues and organs that information has been received and acted upon.

### The cellular basis of feelings and behaviour

Sentience and consciousness have a recognisable cellular basis in humans and other animals. They involve synapses, action potentials, membrane potentials and behavioural synchronisation of a number of neurones (Cook, 2006). The cellular origin of feelings in animals does commence with induced changes in single cells. The route of nervous transmission from sensory system to brain involves two kinds of nerve cell; those that are myelinated and those that are not. Myelin insulates the nerve cell quite effectively from outside interference. The non-myelinated route is thought to be the kind used in the origin of feelings and most of their transmission. Quite clearly such cells are open to environmental influences and it is considered that the membrane potential of any of these is a better indicator of the likely conveyance of information than the action potential itself (Damasio and Carvalho, 2013). Membrane potentials and action potentials figure strongly in plants too.

## THE ‘NERVOUS’ SYSTEM IN PLANTS

One of the primary objections indicated earlier to the possibility for plant sentience was the apparent absence of a nervous system. It is certainly true that the familiar anatomical animal neurone has no equivalent in plants. That was recognised in the 19^th^ century and was also stated by Charles Darwin (1880). He did draw analogies with the behaviour of the root tip (more likely the cap (Trewavas, 2016) and the brain of one of the lower animals. However the lack of obvious neurones does not preclude a functional, excitable but unrecognised, equivalent, capable of electrical transmission which most certainly is present.

J.C Bose, was an Indian scientist, a physicist who worked initially with Rayleigh and was the first to use semiconductor junctions to detect radio signals. He continued research on returning to India in the late 19^th^ century but changed direction and confined himself to investigating the electrophysiology of plant behaviour early in the 20^th^ century. His physical expertise enabled him to construct many pieces of extremely elegant electrical equipment, well before others. He could for example, monitor electrical activity and latent periods of responses within 0.005 seconds. The majority of his studies from this period on, are to be found in some six papers published in the

Proceedings of the Royal Society (to which society he was also subsequently elected in 1920) and one in the Journal of the Linnean Society. There is a further paper in the Royal Society archives which was unpublished, apparently due to objections from Burdon-Sanderson who claimed (wrongly) that only plants with visible movement used action potentials; Bose showed that many others did too. Most of the research material is to be found summarised in two large books, “Plant Response as a Means of Physiological Investigation” (1906) and “Comparative Electrophysiology” (1907). The latter compares plant and animal nervous systems. Additional material is to found in nigh on a dozen books in total. His output was certainly prodigious.

Four years after retirement in 1926, Bose published “The Nervous Mechanism of Plants”. This book describes about one hundred experiments most of whose results are illustrated and contains some additional material not found in the two primary texts. It is not possible to detail all the experiments that Bose performed and instead we have relied on quotations as summary from Bose, (1926).

“The most important fact established in plant response was the nervous character of the impulse transmitted to a distance”. “The conduction of excitation in the plant is fundamentally the same as that of the nerve of the animal”. “The response of the isolated plant nerve is indistinguishable from that of the animal nerve, through a long series of parallel variations of condition” (all page viii). “ In *Mimosa* the velocity of nervous impulse is 400mm/second”.“My recent discovery of the transformation of the afferent or sensory into an efferent or motor impulse in the reflex arc of *Mimosa*, will materially advance our knowledge of the nervous impulse in general” (page ix).

"It has been identified that **excitation is conducted by the phloem of the vascular bundle** and that conduction can be modified experimentally, in the same way as in the animal nerve” (page 215).

Bose established that there was an excitable system in many if not all plants and that it used action potentials. That demonstration, made over 100 years ago, rebuts claims that plants have no ‘nervous’ system, claims that failed to recognise there might be a functional equivalent. But it is the phloem, part of the vascular system, that is conducting electricity over long distances in plants. The phloem has therefore dual functions; transport of organic materials (sugars, amino acids, proteins, peptides, RNA’s of varying size, hormones) as well as conducting electrical impulses and action potentials.

To clarify the distinction with animal neurology, the term phytoneurology and classing individual phloem cells as phytoneurones can be used, when conduction is through excitable phloem cells.

Numerous modern investigations (e.g. Volkov and Ramatunga, 2006; Fromm and Lautner, 2007; Zimmermann et al., 2016; Pickard, 1973 and references therein) have confirmed the validity of these early claims of electrical communication by Bose. Vascular bundles (phloem, cambium and xylem) stretch from the top to the bottom of the individual plant and there is therefore the potential for very long distance communication in large plants and at considerable speeds (Galle et al., 2015; Fromm and Lautner, 2007; Fromm and Bauer, 1994; Yan et al, 2009; Zimmerman et al. 2016). Action potentials in plants can move from 0.5 to 20 cm/sec and the distance covered may be helped by the recently described system potentials (Zimmerman et al., 2016). In addition to action potentials, variation potentials have also been characterized. They are at least 20 fold slower in transmission and may last up to 30 minutes influencing surrounding cell behaviour during this time period. Variation potentials are also dose dependent and more localised near to the site of stimulation.

These two phytoneurological signals (action and variation potentials) rapidly separate from each other following signal initiation. Specific information is thus conveyed by the separation of distance between these two phytoneurological signals as well as amplitude, duration and profile which appear also to be signal specific.

Voltage-gated and mechanosensitive channels are used in the phloem. Instead of initiation by sodium channels in nerves, phytoneurones use chloride efflux throughout specific chloride channels followed by activation of calcium and potassium channels. Plasmodesmata transmit the excitable state and variation potentials to other cells. These phytoneurological signals can be transferred over at least half a metre in young trees and damage or cold shock to one leaf is experienced by other leaves remote from the signal. (Oyarce and Gurovich, 2010; Gurovich and Hermosilla 2009; Lautner et al., 2005)

Action potentials are also accompanied by a cytosolic-free Ca^2+^ wave thus providing excellent cellular interpretation of the information (Choi et al., 2016). Hundreds of downstream proteins (including many sequence-separable calmodulins) in plants are Ca^2+^ sensitive and able to interpret the perceived signal.

### What plant signals induce phytoneurological events and what responses result?

The signals initiating action potentials have been tabulated on several occasions. Numerous aspects of behaviour and development are consequently altered (Fromm and Lautner, 2009; Galle et al., 2015; Pickard, 1973; Trebacz, 1989; Yan et al., 2009). The signals involved are herbivory, (caterpillars, beetles on shoot and nematodes on roots), light-dark changes, temperature variations (cold shock, heat shock), mechanical stimulation, (bending usually resulting from wind impact and inducing stress and strain in cells including and activating mechanosensitive channels in phloem), reductions in mineral uptake, application of saline conditions to roots, watering droughted plants, application of pollen to the stigma, leaf and fruit removal. The consequential changes involve alterations in gene expression, reductions of photosynthesis and increased respiration. Others include a lower rate of phloem translocation, reductions in turgor pressure and stem growth, closure of stomata, induction of hormone synthesis (notably ethylene, abscisic acid, jasmonic acid) and increased nectar secretion. Most of these signals can be classed as indicators or are clearly associated with stressful and potentially damaging conditions.

One critical signal generating action potentials is herbivory. Leaves and stems are damaged and eaten with a high degree of probability by invertebrate and vertebrate herbivores. Below ground, nematodes can create root damage opening up the tissues to fungal infection. As a result of the action potentials generated and variation potentials in cells adjacent to the excitable phloem, defence mechanisms are initiated. One of the commonest reactions is the synthesis and circulation of natural pesticides like caffeine or nicotine, but usually specific to the species. There are an estimated 100,000 natural pesticides reflecting 200 million years of evolutionary arms race between angiosperms and insect pests. Additional responses include wall hardening, the production of gums or attraction of parasitoids as indicated earlier. The herbivore-induced action potential is the first step that leads to a complex set of specific resistance mechanisms that engender memories of differing length.

## THE NERVOUS SYSTEM OF PLANTS CONSISTS OF COMPLEX NETWORKS OF EXCITABLE TISSUES CARRYING ELECTRICAL SIGNALS. IS THIS A BRAIN-LIKE ACTIVITY?

### The need for assessment

Animals are unitary organisms. Rapid movement to find food or avoid being eaten places strong constraints on the phenotype that develops. Assessment is commonly limited to the brain. Memory is accessed and compared to present circumstances activating muscles when needed to initiate movement. Later hormone secretion helps coordinate a whole organism response.

The self-organising plant permits a greater level of distributed control as reflected in plasticity of the phenotype but decisions have to be made after assessment of the prevailing circumstances as well as the requirement to generate the optimal response (Trewavas, 2014). In changing the phenotype resources have to be redirected. This is accomplished by proliferation of new vascular tissue by the cambium to provide these resources to potentially productive branches, by blocking off some vascular elements to those less productive and blocking them entirely to those that provide nothing. Herbivore damage is in part responsible for the need for assessment and specific alteration of phenotypic structure. The cambium itself may be the recipient of the required information from the excitable phloem and the site of assessment here whose need is as important in plants as it is in animals. Without it the phenotype would simply become random and chaotic.

“No form of ganglion however has ever been observed in plants but it is not impossible that the physiological facts may one day receive histological verification” (Bose (1926). page 218). Although Bose recognised the absence of an anatomical animal ganglion in plants, that is, a recognisable conglomerate of nerve cells clustered together, we consider there is one piece of evidence, initially collected by Bose, which may suggest a kind of functional equivalent. It is shown in figure 1 which is taken from figure 54 of Bose (1926). The figure shows one layer of the vascular tissue in *Papaya*. Again quotation reveals Bose assessments. “How reticulated they (vascular bundles) may often be even in the trunk of a tree is shown in the photograph of the distribution of vascular bundles in the main stem of *Papaya* (figure 54). This network of which only a small portion is seen in the photograph girdles the stem throughout its whole length and in this particular case, there were as many as twenty such layers one within the other” (Bose, (1926), page 121). Figure 1 shows the vascular system of this mature plant to consist of numerous vertical vascular elements cross-linked extremely frequently by numerous irregularly-distributed horizontal and tangential connections. A complex network of excitable phloem cells is clearly present.

**Figure 1.**
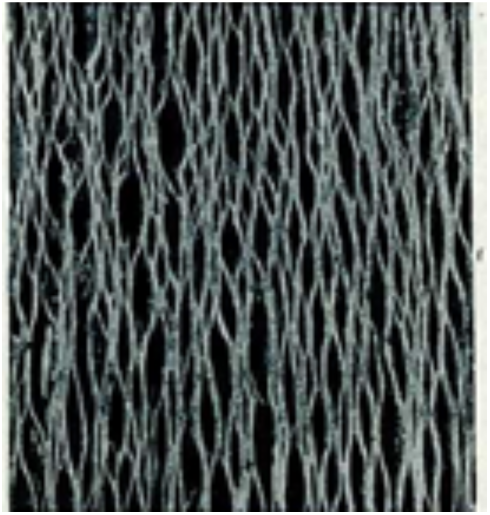
Distribution of vascular tissue in a single stem layer of *Papaya*. There are 20 such layers one inside the other. The bundles are connected through enormous numbers of anastomoses and tangential connections to form a complex excitable structure. “The existence of a system of nerves enables the plant to act as a single organised whole” a requirement perhaps for selection on fitness. Figure and quote taken from figure 54, Bose (1926).

### Leaf excitable phloem networks

The connection with individual leaves is also made by Bose. “The expanded lamina of the leaf in which bundles are spread out in fine ramifications is not merely a specialised structure for the stimulus of light but also a catchment basin for the stimulus of light which is gathered into larger and larger nerve trunks for transmission to the interior of the plant”. And of course other signals too.

There are at least four orders of vein (xylem and phloem) in many angiosperm leaves based on diameter (Sack and Scoffoni, 2013). This hierarchical leaf vein system of angiosperms results in the smallest (finest) veins having a total greater than 80% of the vein length and producing a highly and finely reticulated mesh. There is certainly some degree of irregularity in the fine branching and spatial position.

Leaves generate action potentials in response to cold treatments and mechanical damage from caterpillars and no doubt light and hydraulic signals (Fromm and Bauer, 1994; Zimmermann et al., 2016). Leaves of many species maintain an internal temperature of 21.4 ± 2.2° C throughout the growing season when the external environment varied from 6-30° C, (Helliker and Richter, 2008). A variety of mechanisms (movement into or out of direct sunlight, stomatal aperture, chloroplast movement, hair numbers and reflective /non reflective wax and local leaf number) are used to either warm or cool the leaf keeping the temperature at this homeostat optimum (Trewavas, 2014). Some of these changes take just a few minutes others, a few days. Given the vascular network in the leaf, an action potential generated in part of the leaf would travel throughout. Cells adjacent to the phloem would either experience an action potential themselves or longer lived variation potentials. However an action potential generated in one leaf on a Poplar stem passed into the excitable stem phloem and initiated action potentials further down on leaves remote from the source leaf and on the alternate (acropetal) side (Lautner et al., 2005). The phloem network obviously permits transverse signalling. Bearing in mind the huge numbers of leaves on trees particularly in season, the potential for detailed monitoring of the above ground environment must be enormous. Necessary information on herbivory is likely to be experienced by the whole plant and resistance mechanisms initiated.

### The stem phloem forms an equally complex network

In very young plants, phloem anastomoses (cross links), up to 7000/stem internode in number, have been reported (e.g. Aloni and Sachs, 1973; Aloni and Barnett, 1996). Computer-assisted tomography has been used to identify a complex network of xylem vessels (Brodersen et al., 2011). However xylem does not differentiate in the absence of phloem although the converse is not true (Roberts et al., 1988. Page 47). So the observed vessel network probably ndicates the phloem network too.

In more mature stems and with appearance of additional secondary and supernumery cambia and other features of secondary growth, plant vascular architecture becomes extremely complex. Tangential connections and anastomoses between numerous bundles become very frequent as do radial connections between different stem layers (Carlquist, 1975; Dobbins, 1971; Horak, 1981; Wheat, 1977; Zamski, 1979). These anastomoses do not occur simultaneously in the xylem and phloem and they do construct a “complex netlike structure” already observed in some related 20 families of plants and with further research no doubt most others, confirming Bose observations. The complexity of the excitable phloem network is nothing like the simple structures of vascular tissue presented in text books which are usually limited to seedlings. Woody tissues are penetrated by the phloem and starch deposited which is then mobilised on a seasonal basis.

The majority of dicotyledonous angiosperm species are trees. Very complex networks of phloem can be expected to be present. The assessment capabilities of these excitable phytoneurological networks remain currently unknown but as indicated earlier they should be capable of both local control and control throughout the whole plant since the vascular tissue is present from top of the trunk to the thousands of roots. The key question now: is this network and its behaviour sufficiently complex to be analogous to a mental state?

### Potential behaviour of this complex electrical network?

Networks of all kinds possess emergent properties; properties that originate from the connections between the constituents and nervous systems are one of these (Trewavas, 2007). Even very simple networks of some five interconnected nerve cells exhibit a capability for memory, error correction, time sequence retention and a natural capacity for solving optimisation problems (McCulloch and Pitts, 1943; Hopfield, 1982; Hopfield and Tank, 1986). Learning in nervous systems consists of the construction of either new channels of communication (connections) or altering connection strength between pre-existing neurones (Bray 1990).

Bose (1926) investigated the behaviour of whole vascular tissues which he isolated. He reported that continued stimulation of the phloem phytoneurone increased the size of electrical transmission from the same signal and demonstrated a similar property for the frog nerve. Such capabilities suggest a potential for learning in plants through modification of the phytoneurological connection strength. In this network increased or decreased connection could result from changing the numbers of anastomoses. However testing this potential in the complex mature plant vascular system will not be easy.

Zimmermann et al., (2016) report that there are discrepancies between different publications as to variations of voltage kinetics and magnitudes of action potentials. The information is summarised in a table in their report. While they make some suggestions by way of explanation, there is no recognition that the electrical system is highly branched, that the response is likely holistic and that branching would be variable between individual plants and along single vascular strands (Aloni and Sachs, 1973). While complex branching of the phloem might not explain all such variations it seems to us that it will be a major variable. Measurements need to be conducted with that realisation in mind. Alternatively since the phloem is the primary conductive tissue, electrodes should always be placed into this tissue to try and ensure better uniformity.

### Is this excitable electrical system primed for response and in constant operation?

Both shoots and roots maintain a bioelectric field around themselves (Lund, 1947; McAulay and Scott, 1954; Scott and Martin, 1962). The field has a distinct polarity with different regions exhibiting different potential differences (e.g shoot and root tip more negative than base). These measurements were made on growing organs. The bioelectric fields are in a sense holistic, reflecting the contribution of thousands of cells at any one time. However shoots and roots grow from their respective apices which contain the regions of cell division and growth. The bioelectric field for individual cells must then be dynamic as cells disappear from the zone of division and then subsequently enlarge thus changing their position and their electrical contribution.

However one of the most important observations is that both root and shoot bioelectric fields oscillate by some 30mV in size and with frequencies from 4-15 minutes in roots and 10 to 50 minutes in shoots (Lund, 1947; MacAulay and Scott 1954). Oscillations are usually driven by forms of negative feedback and in this case issuing from the fluctuations of ion flux into and out of cells. A system that oscillates however is primed for response, ready to respond rapidly upon receipt of signals. Measurement of internal electrical potential in tall trees indicate the same pattern of oscillation, or pulsations as Bose (1923) describes them. He was able to demonstrate that these oscillations occur in the endodermis, a group of cells that surrounds the excitable phloem. Later work demonstrated that the endodermis in shoot stems contains the statoliths, the necessary agents detecting gravitational signals (Psaras, 2004; Morita et al., 2002). These observations indicate the electrical system is maintained in an active state and the oscillations keep it in a state of ready alert.

Mechanical, gravitational, electrical signals, temperature and light alter the characteristics of the bioelectric field in the shoot of young plants. When oriented horizontally, the tip/base separation of voltage is now replaced by the equivalent voltage separation across from the upper to the lower stem tissue. The changes in bioelectric field are detectable within one minute. They precede changes in hormone synthesis whose effects on growth are only detectable some 20-30 minutes later (Schrank, 1944; 1945). Recent measurements confirm that electrical changes precede those of growth and may be responsible for them (e.g. Weisenseel and Meyer, 1997; Monshausen et al., 2011).

The root bioelectric field is sensitive to inhibition of growth, temperature osmotic effects and light again. Oscillations in bioelectric field are mimicked by oscillations in ion uptake (Shabala, 2003). The uptake of ions by the root is controlled by the shoot and when dark changes to light an electrical signal provides the necessary information opening up channels and ion movement through the root (Shabala et al., 2007).

## PRIORITISING WHICH SIGNAL TO RESPOND TO

Mental states in animals are thought to be able to prioritise the importance of different signals. Wild plants are subject to huge variations in environmental information from outside and a complex of signals internally. Some of these signals can, when used singly, elicit effective contradictory responses when in combination with some others. Some form of prioritisation of any tissue or organ as to which to respond to first, would then seem essential. From what has been described above some suggestions are now possible.

The set of conditions that initiate action potentials can be loosely grouped as potentially threatening; mechanical damage including predation and physical disruption, cold or heat shock. Physical disruption from bending results from wind signals and those of frictional touch. Changes in tension and compression in all the cells of the tissue can be considerable. The impacts of both wind and touch induce immediate cytosolic Ca^2+^ transients indicative of action potentials (Knight et al., 1993). Bending and flexing in gusty wind can seriously damage leaves and young branches and many plants respond with thickening and increased lignification to reduce movement. We anticipate that excessive movements of leaves will cause extreme flexure in the attaching petiole and the leaf blade and that should generate action potentials along with those generated in the flexing stem or trunk.

However in humans these damaging and temperature treatments are those that deliver pain through nociceptors. By so doing they indicate a priority in both attention and response. The action potentials that are generated in plants will, we suggest, provide a priority to the response against other potential signals. Perhaps the most interesting is how these action potentials are assessed and here the phloem network may be the key. Nothing is known of the behaviour of the phloem anastomoses; will these have any kind of synaptic function? Detailed anatomical and functional analysis of these anastomoses has yet to be performed. Current literature shows almost no awareness of their presence.

The potential for modification of transmission has been referred to earlier which is suggestive at present but no more than that. Networks particularly ones as clearly complex as these should have some potential for assessment, and if not in the phloem itself, then in the cells that surround them and that also experience the specific electrical changes. Signalling in these can induce a variety of molecular changes of differing length and these memories should be accessible through long term modifications in protein expression, particularly chromatin modification.

One other induced action potential that involves the light dark transition may have critical functions in the assessment of shade. The impact is one of reduced nutrition or energy capture unless behaviour is induced to counterbalance. Shade avoidance is a defined syndrome in young plants that attempts to increase shoot growth rates with lower branching rates at the expense of roots and whose function is to overgrow the competition. A daily assessment at the light/dark transition may be the means of making that assessment although in large woody angiosperms likely complex.

Signals that do not induce action potentials seem at present to be most notably those of gravity. In green stems, the statoliths enabling gravitropic responses are located in the endodermis, a group of cells surrounding the excitable phloem (Psaras, 2004; Morita et al., 2002). But if green plants grown in pots are inverted over a light source the gravity response is overridden. Phytochrome A, a light sensitive pigment is found at highest concentrations in these endodermal cells too (Hisada et al., 2000). In this case the prioritisation might simply be brute force in the responsive cells with stronger promoters for light reactions against those for gravitropism responses. Alternatively in the root cap which normally contain cells with statoliths, placement of other signals at right angles to gravity leads to loss of the statoliths (Massa and Gilroy, 2003; Eapen et al., 2005). In this case priority is gained by elimination of alternative sensing.

If a plant is subject to shade situations and to a mild deprivation of water, which response would be priority? The shade avoidance syndrome normally leads to enhancement of stem growth at the expense of the root. Water deprivation is normally claimed to lead to reductions in shoot growth, enhancement of root growth and if necessary loss of leaves. Would the stem grow faster or resources instead be given to enhance root exploration for water? Would the phytoneurological network indicated above resolve such situations and thus provide a way in which the individual plant can assess the environmental situation and determine which physiological group is pre-eminent? These questions need better resolution if understanding of the behaviour of wild plants and trees is to be gained.

## CONCLUSION

The reticulated excitable phloem system described above offers a potential for assessment of signals and perhaps their prioritisation. The phytoneurological system is present throughout any growing plant and thus should be capable of dealing with local signals as well as those that require a more integrated response. Both local and long distance changes are characteristic of higher plants. Bose (1926) suggests that it provides for the construction of an integrated whole organism. The vascular network is some kind of complex interactive system and once stimulated has the potential for assessment through possible feedbacks and alterations of connection strength. Whether it should be regarded as a functional equivalent to a fairly primitive brain cannot be determined until its properties are more clearly defined by research. But a feature of most plants is phenotypic plasticity and any kind of phytoneurological system has to accommodate that too. As in other organisms it is no doubt the mixture of chemical and nervous connections that is used for communication throughout the organism.

This article commenced by pointing out that lack of obvious movement in plants has led to a downgrading of any kind of nervous control altogether and this needs rebalancing. The article here raises important issues that have been neglected and that require suitable answers not least from electrophysiologists of all kinds. With recognition that this nervous system might act holistically, some issues that have dogged this area of research might be better understood.

## Acknowledgments

This research was supported by Spanish Ministry of Education, Culture and Sport through a “Stays of professors and senior researchers in foreign centres” fellowship to P.C.

